# Recurrent selection with glufosinate at low rates reduces the susceptibility of a *Lolium perenne* ssp. *multiflorum* population to glufosinate

**DOI:** 10.1101/2020.07.04.182733

**Authors:** Maor Matzrafi, Sarah Morran, Marie Jasieniuk

## Abstract

Repeated applications of herbicides at the labelled rates have often resulted in the selection and evolution of herbicide-resistant weeds capable of surviving the labelled and higher rates in subsequent generations. However, the evolutionary outcomes of recurrent herbicide selection at low rates are far less understood. In this study of an herbicide-susceptible population of *Lolium perenne* ssp. *multiflorum*, we assessed the potential for low glufosinate rates to select for reduced susceptibility to the herbicide, and cross-resistance to herbicides with other modes of action. Reduced susceptibility to glufosinate was detected in progeny in comparison with the parental population following three rounds of selection at low glufosinate rates. Differences were mainly observed at the 0.5X, 0.75X, and 1X rates. Comparing the parental susceptible population and progeny from the second and third selection cycle, the percentage of surviving plants increased to values of LD_50_ (1.31 and 1.16, respectively) and LD_90_ (1.36 and 1.26, respectively). When treated with three alternative herbicides (glyphosate, paraquat, and sethoxydim), no plants of either the parental or successive progeny populations survived treatment with 0.75X or higher rates of these herbicides. The results of this study provide clear evidence that reduced susceptibility to glufosinate can evolve in weed populations following repeated applications of glufosinate at low herbicide rates. However, the magnitude of increases in resistance levels over three generations of recurrent low-rate glufosinate selection observed is relatively low compared with higher levels of resistance observed in response to low-rate selection with other herbicides (three fold and more).

## 1. INTRODUCTION

Weeds are the major pests limiting crop production in agricultural systems.^1^ Treatment with herbicides is by far the most effective method of controlling weeds although repeated applications of herbicides select for, and can result in, the evolution of herbicide resistance.^2,3^ When applied at labelled field rates, herbicides have often selectively favoured individuals possessing major resistance alleles and target-site resistance (TSR) that spread rapidly within and among weed populations.^2,3^ However, the evolutionary outcomes of recurrent herbicide selection at rates lower than the labelled rates are far less understood. It has been suggested that repeated applications of herbicides at lower than labelled rates selects for polygenic herbicide resistance in weeds.^4,5^ Consequently, each subsequent generation of selection is predicted to result in a slow shift of the entire population towards resistance. For instance, recurrent selection with low rates of dicamba^6^ and glyphosate^7^ only resulted in 2.15- to 3-fold lower susceptibility of *Amaranthus palmeri* to these herbicides over three to four generations in comparison to the parental populations. In contrast, however, three generations of recurrent selection with low rates of diclofop-methyl resulted in high level (56-fold) of resistance in *Lolium perenne* ssp. *rigidum*.^8^

Recurrent selection at low herbicide rates has sometimes also resulted in the selection of progeny with cross-resistance to other herbicides. For instance, repeated applications of pyroxasulfone at low rates resulted in the selection of a *L. perenne* ssp. *rigidum* population that was resistant to pyroxasulfone and cross-resistant to chlorsulfuron, diclofop-methyl and S-metolachlor.^9^ In *Avena fatua*, repeated applications of diclofop-methyl, an ACCase (acetyl CoA carboxylase)-inhibiting herbicide, at low rates for three consecutive generations resulted in the selection of progeny populations with reduced susceptibility to diclofop-methyl and cross-resistance to ALS-inhibiting herbicides.^10^

*L. perenne* ssp. *multiflorum* (Italian ryegrass) is one of the major weed species in orchards, vineyards, field crops, and fallow fields of California.^11,12^ Extensive herbicide use has exerted strong selection that has resulted in the evolution of herbicide resistance in many populations of this weed species in California.^12–16^ Resistance to glyphosate,^12,14,17^ paraquat, and the ACCase inhibitor, sethoxydim,^15^ as well as multiple herbicide resistance to these three herbicides plus acetolactate synthase (ALS) inhibitors^15,16^ have been confirmed in populations across the agricultural landscape of northern California. Consequently, the management of herbicide-resistant *L. perenne* ssp. *multiflorum* has become a major challenge in California annual and perennial cropping systems.

Glufosinate is an alternative non-selective post-emergence herbicide that can still be used to control herbicide-susceptible and most herbicide-resistant *L. perenne* ssp. *multiflorum* in California as only two populations with glufosinate resistance have been documented to date.^14^ Both are populations with low resistance levels (1.6-2 fold) compared to the standard susceptible population. Worldwide, six additional cases of glufosinate resistance have been reported in *Lolium* species.^18^ In Oregon, both target site and non-target site mechanisms were suggested as endowing resistance to glufosinate in *L. perenne* ssp. *multiflorum* populations.^19,20^

The relatively high cost of glufosinate, as well as the increasing abundance of weeds resistant to alternative herbicides, may drive farmers to apply more glufosinate, but at reduced rates. This, among other drivers such as herbicide applications at non-optimal weed size, inappropriate weather conditions, and insufficient spray coverage may result in sublethal rate herbicide selection. Thus, there is a need to assess the potential for recurrent selection with glufosinate at low rates in *L. perenne* ssp. *multiflorum*, the weed species with a high propensity to evolve resistance to herbicides with different modes of action.

Hence, the objectives of the present study were: (1) to evaluate the potential for low glufosinate rates to select for reduced susceptibility to the herbicide and (2) to determine if selected populations are cross-resistant to herbicides with other modes of action that are commonly used to control *L. perenne* ssp. *multiflorum* in orchards and vineyards of California.

## 2. MATERIALS AND METHODS

### 2.1 Plant material

Seeds of a previously characterized herbicide-susceptible population of *L. perenne* ssp. *multiflorum* from a vineyard in Sonoma County, California^15,16^ constituted the parental population (P_0_) for this study. Seeds were germinated on moistened filter paper in Petri plates with 1% v/v Captan 80 WDG (Agri Star, Ankeny, IA, USA) and incubated at ambient temperature under a 12-h photoperiod provided by fluorescent lights (160 µmol m^2^ s^-1^). Seedlings at the one- to two-leaf stage were transplanted into plastic pots (5 cm height × 4.5 cm diameter) filled with UCD Ron’s soil mix (1:1:1:3 sand/compost/peat/dolomite). Pots were maintained in a growth chamber (model PGV 36; Conviron Ltd., Winnipeg, MB, Canada) under 25/19 + 3° C (day/night) temperature and 12-h photoperiod using high pressure sodium lamps (600 µmol m^2^ s^-1^).

### 2.2 Recurrent selection with glufosinate at low rates

Six hundred P_0_ seedlings at the three- to four-leaf stage (8-10 cm tall) were divided into three sets of 200 seedlings. Each set of 200 P_0_ seedlings was treated with glufosinate (Rely 280®, Bayer CropScience) at one of three rates (123, 246, or 492 g ai ha^−1^), equivalent to 0.125X, 0.25X, and 0.5X of the labelled field rate (984 g ai h^-1^). Glufosinate was applied using an automated track sprayer equipped with a 8001E flat-fan nozzle (TeeJet Technologies, Springfield, IL, USA) calibrated to deliver 187 L ha^-1^ at 296 KPa. Treated plants were maintained in the growth chamber under the conditions described above. The number of surviving plants was recorded 21 days after treatment (DAT). Glufosinate at the 492 g ai ha ^-1^ resulted in highest plant mortality (76.5%) among the rates used. All 47 surviving plants were transplanted into larger round plastic pots (2.37 L) filled with commercial potting mix (LC1, Sun Gro Horticulture, AB, Canada), grown to maturity under the conditions described above, and allowed to cross-pollinate. Mature seeds that were collected from these plants, designated the P_1_ generation, were air-dried at room temperature and stored at 4° C for four to six weeks to overcome dormancy and maximize germination for a subsequent round of selection.

For the second round of selection, P_1_ seeds were germinated and seedlings transplanted into pots and grown to the three- to four-leaf stage, as described above. For this selection round, 900 P_1_ seedlings were divided into three sets of 300 seedlings. Each set of 300 P_1_ plants was treated with glufosinate at one of three slightly higher rates of glufosinate (0.5X, 0.75X, and 1X) than in the first round of selection. Approximately 50 surviving plants were selected from the 738 g ai ha-1 rate (0.75X), which resulted in 79% plant mortality, and transplanted into larger pots, grown to maturity, and allowed to cross-pollinate. Mature seeds were collected, designated the P_2_ generation, and stored for four to six weeks to overcome dormancy. A similar approach was taken for an additional round of selection with glufosinate at three higher rates, equivalent to 0.75X, 1X, and 1.25X the labelled field rate, to produce the P_3_ generation of seeds.

### 2.3 Dose-response of the parental and selected populations to glufosinate

To compare the response of the parental population (P_0_) and the selected progeny populations (P_1_, P_2_, P_3_) to glufosinate, seedlings at the 3- to 4-leaf stage (8-10 cm) from each generation were treated with glufosinate at seven rates (0.125X, 0.25X, 0.5X, 0.75X, 1X, 2X and 4X). Following treatment, plants were kept for 21days in the growth chamber under the environmental conditions described earlier. The experiment was conducted in a complete randomized design (CRD) with 10-12 replications of individual plants from each generation per treatment and repeated. Plant survival was recorded 21 DAT.

### 2.4 Cross-resistance to glyphosate, paraquat, and sethoxydim

In an experimental design (CRD) similar to that described above for the glufosinate dose-response study, cross-resistance to other herbicides was assessed for the parental population (P_0_) and for the three selected progeny populations (P_1_, P_2_, P_3_). The experiment was repeated. Seedlings at the 3- to 4-leaf stage (8-10 cm) from each generation were treated with seven rates (0.125X, 0.25X, 0.5X, 0.75X, 1X, 2X and 4X) of glyphosate (Roundup PowerMax^®^, Monsanto; 1X = 867 g ae ha^-1^), paraquat (Gramoxone SL 2.0^®^, Syngenta Crop Protection; 1X = 560 g ai ha^-1^) and sethoxydim (Poast^®^, BASF Corporation; 1X = 515 g ai ha^-1^). Crop oil concentrate (COC; Helena Chemical Company, Collierville, TN) at 1% V/V and Nonionic surfactant (NIS; Helena Chemical Company) at 0.25% V/V were added to spray solutions containing sethoxydim and paraquat respectively. Treated plants were kept in a growth chamber under the environmental conditions described earlier and plant survival recorded 21 DAT.

### 2.5 Statistical analyses

Plant survival and shoot biomass data were pooled over the two runs of each experiment due to nonsignificant differences between runs for all experiments. For all herbicides, plant survival data from the dose-response experiments for the P_0_, P_1_, P_2_, and P_3_ populations were fit to a binomial two-parameter log-logistic model using the drc package of R version 3.5.1^21^ and the LD_50_ values (rate required for 50% plant mortality) and LD_90_ values (rate required for 90% plant mortality) estimated.

To further assess cross-resistance of the P_0_, P_1_, P_2_, and P_3_ populations to glyphosate, paraquat, and sethoxydim, data on the percentage of fresh shoot weight reduction from the dose-response experiments were fit to a nonlinear sigmoidal logistic three-parameter model.^22^

All data was visualized using SigmaPlot (ver. 13) software (Systat Software Inc., San Jose, CA, USA).

## 3. RESULTS AND DISCUSSION

### 3.1 Recurrent selection with glufosinate at low rates

As expected, the percentage of P_0_ plants surviving treatment with glufosinate was inversely related to the herbicide rate, with 100%, 90.5%, and 23.5% of plants surviving 0.125X, 0.25X, and 0.5X times the labelled field rate of glufosinate, respectively, 21 DAT (Table 1). Repeated selection with glufosinate at low rates over three consecutive generations produced three successive populations (P_1_, P_2_, and P_3_) of progeny with an increasing percentage of plants surviving treatment with glufosinate at a specific rate. Thus, whereas only 23.5% of P_0_ plants survived the 0.5X rate of glufosinate, a larger percentage (71%) of P_1_ plants survived the same dose in the next generation (Table 1). Similarly, only 21% and 5% of P_1_ plants, but 33% and 12% of P_2_ plants, survived the 0.75X and 1X rates of the herbicide, respectively, indicating that selection with glufosinate at low rates had reduced susceptibility to the herbicide, as assessed by the increasing proportions of survivors at each rate over generations.

**Table 1.**
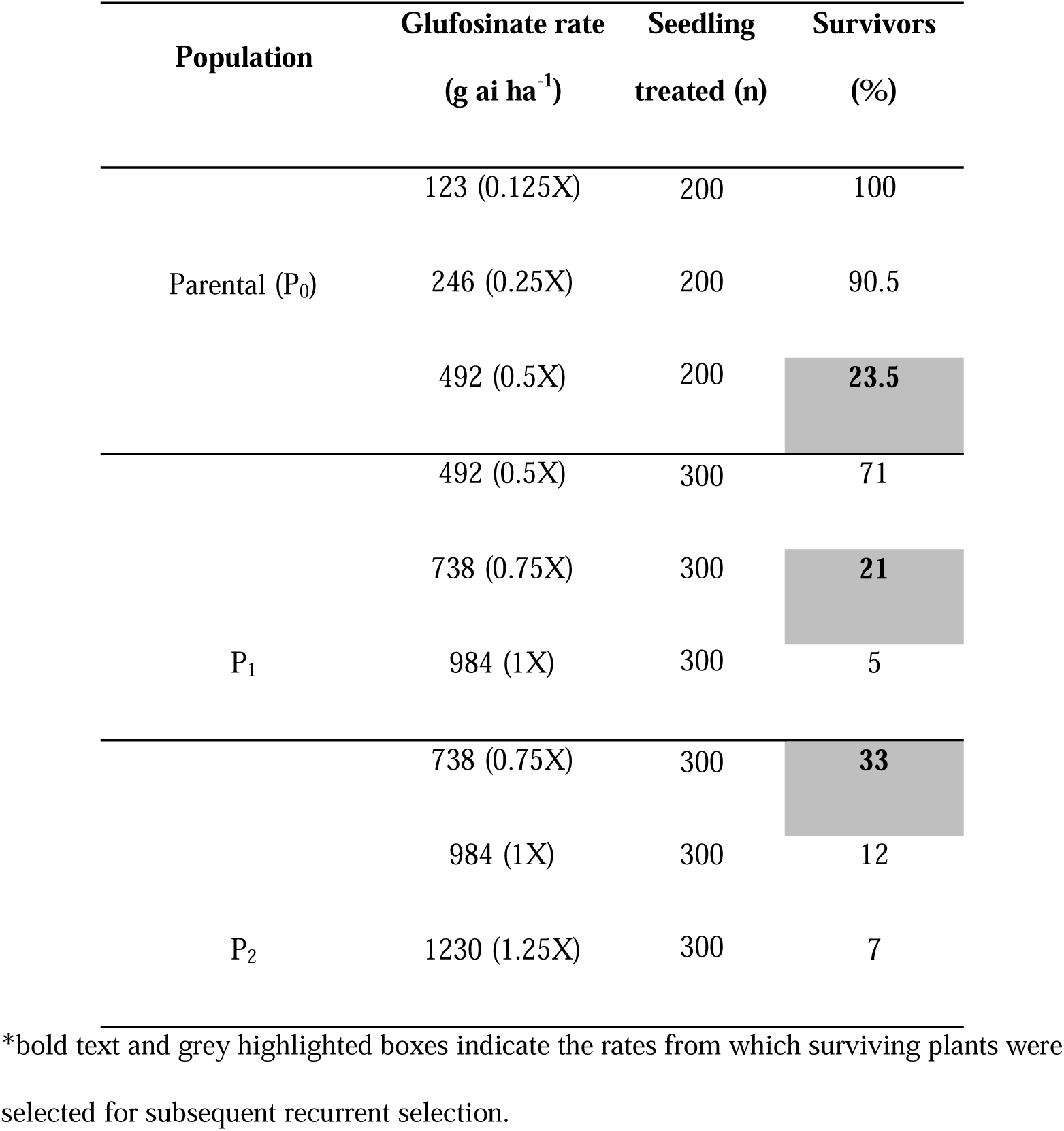
Percentage of *L. perenne* ssp. *multiflorum* plants surviving treatment with glufosinate at low (i.e., lower than the recommended labelled) rates.

### 3.2 Dose-response of parental and selected populations to glufosinate and other herbicides

Progeny populations P_2_ and P_3_ exhibited reduced susceptibility to glufosinate at rates ranging from the 0.5X to 1X the labelled field rate in comparison with the parental population (P_0_) (Fig. 1). LD_50_ and LD_90_ values for populations P_2_ (592.08 and 1117.58 g ai/ae ha^-1^, respectively) and P_3_ (529.2 and 1038.16 g ai/ae ha^-1^, respectively) were higher in comparison with those of the P_0_ (452.39 and 816.66 g ai/ae ha^-1^, respectively) and P_1_ (429.95 and 888.82 g ai/ae ha^-1^, respectively) populations (Table 2). The level of resistance, as measured by the Resistance Index (RI) calculated using LD_50_ values and the parental population P_0_ as the susceptible standard, revealed RI values of 0.95, 1.31, and 1.16 for the P_1_, P_2_, and P_3_ populations, respectively. Based on LD_90_ values, RI values were 1.08, 1.36, and 1.26 for the P_1_, P_2_, and P_3_ populations, respectively. Our results clearly show that the percentage of plants surviving glufosinate was higher for the P_2_ and P_3_ progeny populations compared to the parental population P_0_ (Table 2), however, in comparison to low-rate selection studies with other herbicides, the level of resistance did not increase substantially.

**Table 2.** Parameter estimates and associated model statistics for the log-logistic dose-response curves of plant survival following treatment with glufosinate at low doses.

| **Population** | <sup>a</sup>LD<sub>50</sub><br>(g ai/ae ha<sup>-1</sup>) | **SE** | **RI<sup>d</sup>**<br>(P<sub>n</sub>/P<sub>0</sub>) |
| --- | --- | --- | --- |
| P<sub>0</sub> | 452.39 (382.77-522.02)<sup>c</sup> | 35.52 | -- |
| P<sub>1</sub> | 429.95 (351.98-507.93) | 39.78 | 0.95 |
| P<sub>2</sub> | 592.08 (502.46-681.7) | 45.72 | 1.31 |
| P<sub>3</sub> | 529.2 (444.75-613.65) | 43.08 | 1.16 |
|  | <sup>b</sup>LD<sub>90</sub><br>(g ai/ae ha<sup>-1</sup>) | **SE** | **RI**<br>(P<sub>n</sub>/P<sub>0</sub>) |
| P<sub>0</sub> | 817.66 (640.4-994.92) | 90.44 | -- |
| P<sub>1</sub> | 888.82 (656.12-1121.5) | 118.72 | 1.08 |
| P<sub>2</sub> | 1117.58 (839.05-1396.1) | 142.10 | 1.36 |
| P<sub>3</sub> | 1038.16 (782.07-1294.3) | 130.66 | 1.26 |

<sup>a</sup>LD<sub>50</sub> - represents the rate that results in 50% mortality.
<sup>b</sup>LD<sub>90</sub> - represents the rate that results in 90% mortality.
<sup>c</sup>Values in parentheses indicate 95% confidence intervals.
<sup>d</sup>RI, the Resistance Index, is a population's LD<sub>50</sub> or LD<sub>90</sub> divided by the value of the same parameter for the parental population, P<sub>0</sub>.

**Figure 1.**
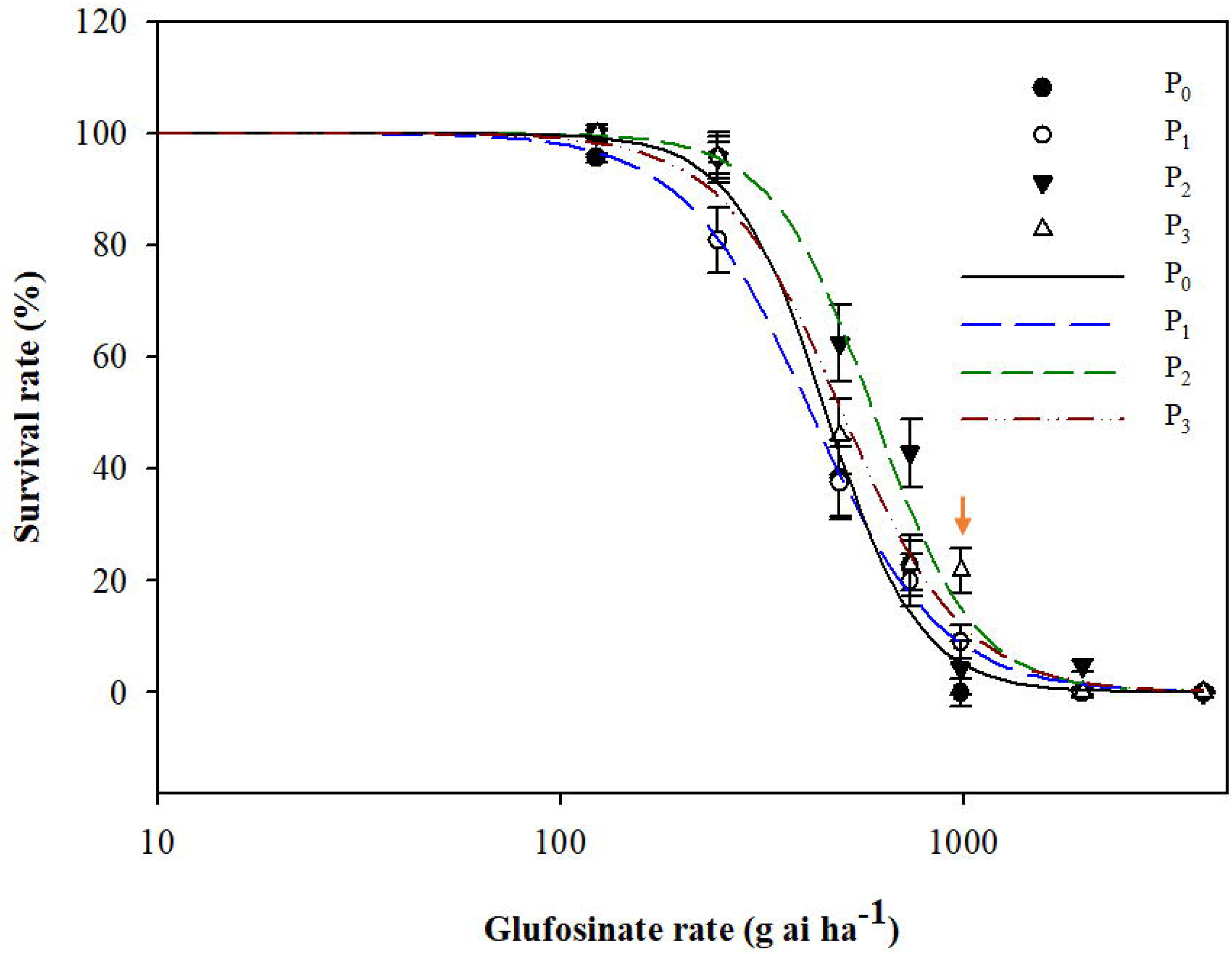
Dose-response of *L. perenne* ssp. *multiflorum* parental (P_0_) and three successive generations (P_1_, P_2_, P_3_) of progeny, selected with low glufosinate rates, to treatment with glufosinate in the greenhouse. Lines are the predicted values for percentage survival. Red arrow indicates the labelled field rate (984 g ai h^-1^).

### 3.3 Cross-resistance to glyphosate, paraquat, and sethoxydim

Interestingly, no plants of the parental population or the P_1_, P_2_, and P_3_ progeny survived glyphosate, paraquat, and sethoxydim treatments equal to and greater than 0.75X the labelled rates of these herbicides (Fig. 2A-C, Table 3). Busi et al.^5^ suggested that selection using low rates may hasten the evolution of polygenic herbicide resistance, especially in cross-pollinated species such as *L. perenne* ssp. *multiflorum*. The authors suggest that reduced sensitivity to chlorsulfuron was observed in progeny from low-rate diclofop methyl selection, apparently as a result of enhanced detoxification of both herbicides.^23^ Cross-resistance to glufosinate and glyphosate was previously suggested in *L. perenne* from Oregon and the resistance hypothesized to be non-target-site related.^20^ In this study, reduced susceptibility to the 0.5X rate of glyphosate was detected following two and three generations of selection with low rates of glufosinate (Table 3) but further research is required to determine whether this is due to cross-resistance.

**Table 3.** Percentage of plants from the parental population (P_0_) and three generations of progeny (P_1_-P_3_) that survived treatment with glyphosate, paraquat, and sethoxydim 21 DAT^a^.

|  |  | % survivors at each herbicide rate |  |  |  |  |  |  |  |
| --- | --- | --- | --- | --- | --- | --- | --- | --- | --- |
|  | Population | 0 | 0.125 | 0.25 | 0.5 | 0.75 | 1 | 2 | 4 |
| Glyphosate | $P_0$ | 100 | 100 | 45 | 0 | 0 | 0 | 0 | 0 |
| | $P_1$ | 100 | 100 | 20 | 0 | 0 | 0 | 0 | 0 |
| | $P_2$ | 100 | 100 | 20 | 20 | 0 | 0 | 0 | 0 |
| | $P_3$ | 100 | 80 | 20 | 20 | 0 | 0 | 0 | 0 |
| Paraquat | $P_0$ | 100 | 0 | 0 | 0 | 0 | 0 | 0 | 0 |
| | $P_1$ | 100 | 10 | 0 | 0 | 0 | 0 | 0 | 0 |
| | $P_2$ | 100 | 0 | 0 | 0 | 0 | 0 | 0 | 0 |
| | $P_3$ | 100 | 20 | 0 | 0 | 0 | 0 | 0 | 0 |
| Sethoxydim | $P_0$ | 100 | 10 | 10 | 0 | 0 | 0 | 0 | 0 |
| | $P_1$ | 100 | 0 | 0 | 0 | 0 | 0 | 0 | 0 |
| | $P_2$ | 100 | 0 | 0 | 0 | 0 | 0 | 0 | 0 |
| | $P_3$ | 100 | 10 | 0 | 0 | 0 | 0 | 0 | 0 |

<sup>a</sup> $n = 12$ for all other herbicides.

**Figure 2.**
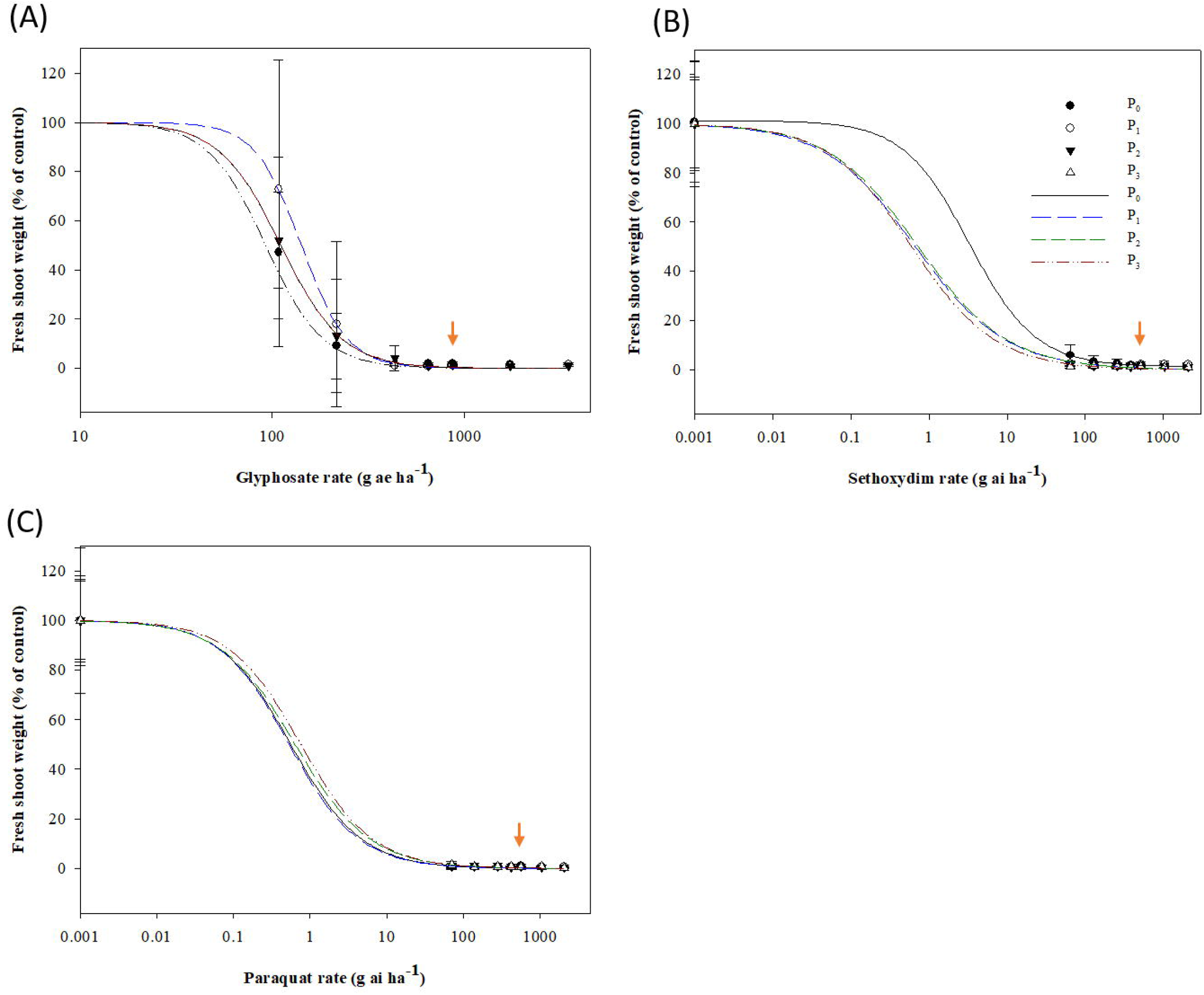
Dose-response of *L. perenne* ssp. *multiflorum* parental (P_0_) and three successive generations (P_1_, P_2_, P_3_) of progeny, selected with low rates of glufosinate, to treatment with glyphosate (A), sethoxydim (B) and paraquat in the greenhouse. Lines are the predicted values for fresh shoot weight. Red arrows indicate the labelled field rates for glyphosate (867 g ae ha^-1^), paraquat (560 g ai ha^-1^) and sethoxydim (515 g ai ha^-1^).

In most recurrent low rate selection studies, resistance level exceeded 3-fold after three generations of low-rate selection.^6,8,10,24^ In this study, the magnitude of increases in resistance levels over three generations of recurrent low-rate glufosinate selection observed contrast with the higher levels of resistance observed in response to low-rate selection with other herbicides. However, the results are consistent with previous studies of glufosinate resistance in *Lolium* species, which generally observe lower levels of resistance to glufosinate with R/S ratios ranging from 1.6 to 2.8 fold.^14,19,20^ Our earlier work^14^ also found significant variability in response to glufosinate among individuals in California populations of *L. perenne* ssp. *multiflorum* and a strong influence of environmental conditions on glufosinate efficacy and sensitivity, which has also been detected in *Raphanus raphanistrum*^25^ and *A. rudis, A. palmeri* and *A. retroflexus*.^26^ Whether the evolution of glufosinate resistance in weed populations is more complex than resistance evolution to other herbicides remains to be investigated. However, the results of this study provide clear evidence that reduced susceptibility to glufosinate can evolve in weed populations following repeated applications of glufosinate at low herbicide rates.

In summary, in this study we showed that three generations of recurrent selection with glufosinate at low rates (i.e., lower than the labelled field rates) was sufficient to reduce the susceptibility of subsequent generations (P_1_-P_3_) of progeny to the herbicide compared with the parental population (P_0_) (Fig. 1). Our findings are consistent with the results of other studies showing that recurrent low-rate selection may lead to the evolution of herbicide resistance.^5,6,10,24^ Reduced susceptibility to paraquat and sethoxydim with successive generations of glufosinate selection was not observed. The increases in frequency of plants surviving increasing glufosinate rates each successive generation may reflect a shift in mean response at the population level indicative of directional selection on quantitative trait variation and, possibly, non-target site related glufosinate resistance.

## ACKNOWLEDGMENTS

We are grateful to Chad Fautt for greenhouse and laboratory assistance, and Hannah Clifton for greenhouse maintenance. This study was funded by Bayer CropScience AG and by BASF Corporation. No conflicts of interest have been declared.

